# DrawAlignR: An interactive tool for across run chromatogram alignment visualization

**DOI:** 10.1101/2020.01.16.909143

**Authors:** Shubham Gupta, Justin Sing, Arshia Mahmoodi, Hannes Röst

## Abstract

Multi-run alignment is widely used in proteomics to establish analyte correspondence across runs. Generally alignment algorithms return a cumulative score, which may not be easily interpretable for each peptide. Here we present a novel tool, DrawAlignR, to visualize each chromatographic alignment for DIA/SWATH data. Furthermore, we have developed a novel C++ based implementation of raw chromatogram alignment which is 35 times faster than the previously published algorithm. This not only enables users to plot alignment interactively by DrawAlignR, but also allows other software platforms to use the algorithm. DrawAlignR is an open-source web application using R Shiny that can be hosted using the source-code available at https://github.com/Roestlab/DrawAlignR.

## Introduction

High-throughput tandem mass spectrometry, specifically data-independent acquisition (DIA), is a deterministic and reproducible approach to studying the proteome of many biological samples [1]. The resulting, highly multiplexed DIA data is often analyzed using a peptide-centric approach, which uses targeted libraries to query the raw spectra. These library-based workflows extract MS2 ion chromatograms and subsequently extract features from it [2, 3, 5, 14]. Since features are picked independently across multiple runs, this introduces inconsistency in peptide quantification, leading to missing quantitative values in the final data matrix [15]. To establish consistency and reduce the number of missing values, retention-time alignment is performed to match peptide quantification across multiple runs [4]. Considering the potentially strong effect of (mis-)alignment on a peptide data-matrix, it is important to validate the alignment visually. Although there are tools available to visualize raw chromatogram and feature-selection [5, 6,16, 17], there is a lack of open-source tools to visualize chromatographic alignment across multiple DIA-runs.

We have previously published an R-package, DIAlignR, that uses raw MS2 extracted-ion chromatograms (XICs) together with high-scoring features for hybrid RT-alignment, which outperforms previous approaches in the alignment of heterogeneous runs acquired over long time periods [7]. However, this implementation was slow, did not provide interactive visualization of the alignment and any visualization required considerable effort and expert knowledge in R scripting. In this paper we present a modern C++ 11 software library that provides a fast re-implementation of the alignment algorithm and an interactive visualization tool, DrawAlignR, to visualize the alignment of chromatograms. The visualization is implemented as an R Shiny app [8] which uses standard open-source formats such as mzML and sqMass (chromatogram files) and tsv and osw (OpenSWATH results files).

## Implementation

Pairwise alignment of extracted-ion chromatograms (XICs) starts with intensity and retention time extraction followed by time-series smoothing, which in total has a computational cost of *θ*(n). Subsequent steps performed in DIAlignR [7] are:1) computing a similarity matrix, 2) constraining similarity with a global fit and 3) performing dynamic programming for affine alignment. These algorithms are computationally expensive, with *θ*(n^2^) complexity in memory and computation time (Suppl Note 1). Since XICs usually are small in length (200-300 data points), the memory requirements are relatively modest. However, execution time was substantial due to the use of non-optimized R code. To decrease execution time, we ported these quadratic functions to C++ libraries (Fig. 1*a*).

**Figure 1:**
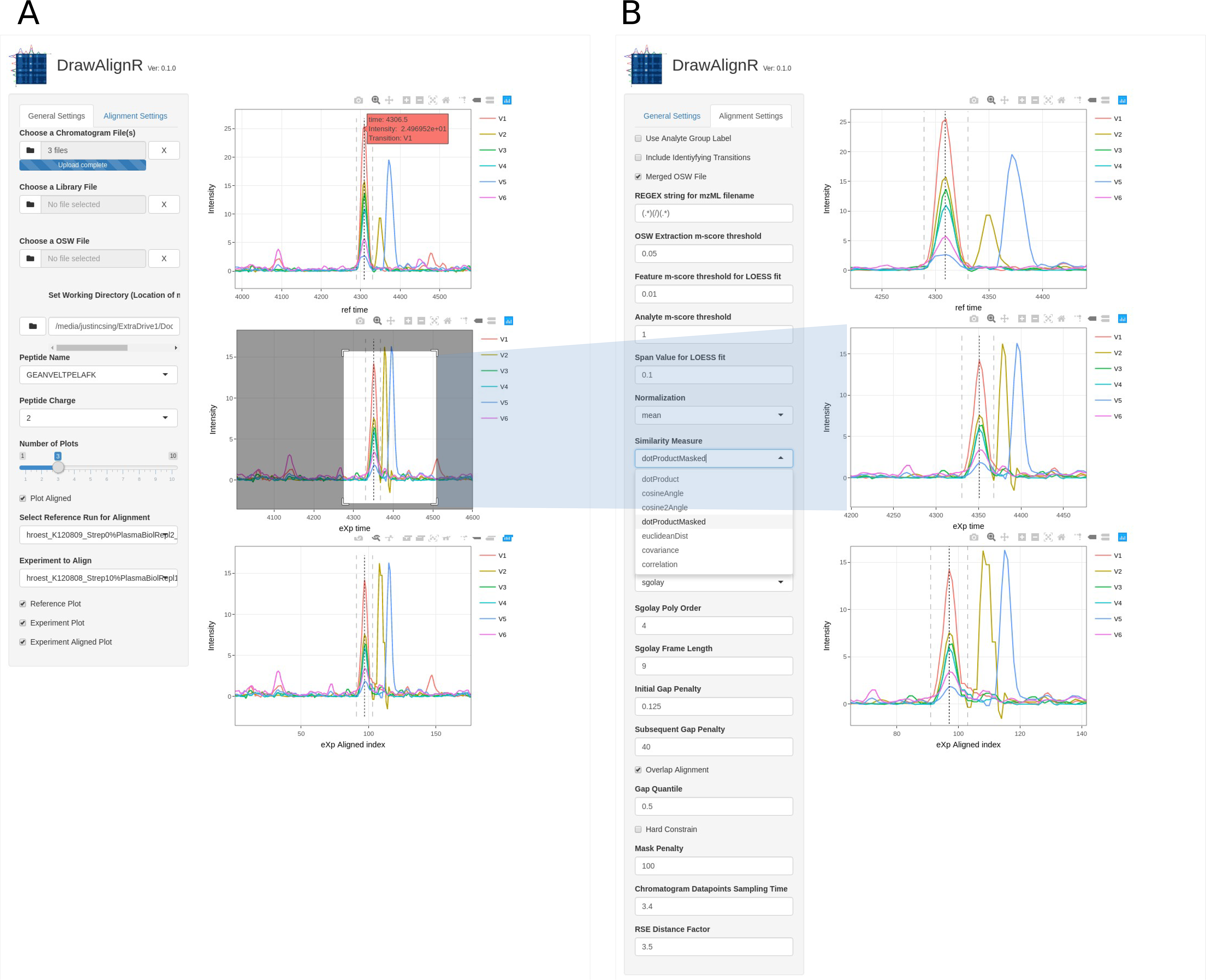
Alignment algorithm in C++. a) We provide a novel C++ library with highly efficient implementations of the raw data alignment functionality of DIAlignR. The library uses modern C++11 and is based on object-oriented C++ with heavy use of STL data structures and algorithms. Abstraction layers allow users to integrate this code easily with R and Python. b) Execution time for alignment of 1000 peptides using Cpp library and R code (Also see Suppl Note 1). The R code was implemented in DIAlignR version 0.1.0, C++ libraries have been made available with the latest release.

### Similarity matrix calculation and constraining

We provide a modern C++ 11 implementation that uses STL algorithms and data structures throughout to compute various similarity matrices (e.g. cosine, dot-product, euclidean distance etc.) directly from raw data XICs. Our use of extensive unit testing and re-use of existing algorithms results in concise and clear code.

### Affine alignment

We provide an object-oriented, modern C++ implementation using the object `AffineAlignObj` which contains the required matrices M, A, B, traceback and alignment path from dynamic programming [7]. We have optimized memory management, allowing users to re-use the previously allocated memory in a transparent fashion, without requiring user intervention.

We have taken advantage of the **Rcpp** package to provide an interface of our C++ code directly to the R programming language (Fig. 1*a*) [9, 10]. This package first creates `RcppExports.cpp` containing functions with wrapped C++ functions from `Rmain.cpp` (Suppl Note 2). These functions are suitable for being called by R using `.Call` (in RcppExports.R) that provides interface for data-interchange between R and C++.

We used Cython (C-Extension for Python) to wrap the C++ library and make it accessible to Python. For this, a.pxd file was created that contains only the functions and class declarations from the C++ library. A companion.pyx file was then written that creates the actual wrapper code in Cython and which is translated by the Cython compiler to generate a.cpp file. A C++ compiler can then create the required shared object for the Python module DIAlignPy.

DrawAlignR has been implemented in R to develop an interactive user interface with rich graphics for visualizing chromatograms, and across-runs peak alignment. The tool was built using R’s Shiny package, which allows easy development of interactive tools accessible via a web browser [8]. Extracted-ion chromatographic data is extracted through R’s mzR package, which retrieves intensity and retention time values from standardized mzML file formats. An mzR object containing pointers mapping fragment ion ids to their respective indexed chromatographic data position in the mzML file is cached into memory for efficient data retrieval. In addition, DrawAlignR is also capable of reading the more efficient sqMass chromatogram file formats using PyMSNumpress and the reticulate R-Python interface. Whenever a peptide is selected from a list of peptides, extracted from either a user supplied library file or an osw results file, an extracted-ion chromatogram is automatically retrieved for the given peptide for each chromatogram run file is supplied. If more than one chromatogram run file is supplied, there is the option of visualizing the pairwise alignment of peptides across runs using DIAlignR, where the alignment is visualized as the shifted RT of an experiment run in correspondence to the selected reference run.

## Results

Our highly efficient C++ library performs alignment in an order of magnitude faster than previous R based algorithms. For a peptide with 10 minute wide XICs (180 data points), C++ based alignment takes only 3.5 ms compared to 116 ms with R code (Figure 1*b*). This 35-fold increase in performance is consistently observed for different chromatogram lengths, thus, each computation required for alignment is inherently slower in R. This means that alignment of whole experiments are now feasible with the efficient C++ implementation, processing up to 300 peptides per second and completing alignment of a moderately large LC-MS/MS experiment with 10,000 peptide precursors in less than one minute.

Our newly developed open source C++ library is platform independent and can be interfaced with multiple (scripting) languages. We have additionally written an abstraction layer for the Python programming language (Figure 1*a*), which is increasingly popular in the proteomics field [11, 18, 19]. This makes it possible to integrate MS2 chromatogram based alignment with existing tools eg. pyopenms [11], TRIC [12] etc.

DrawAlignR is a novel tool to visualize the alignment of XICs interactively, which is not possible with currently available visualization software. This allows users to experiment with chromatographic alignment on their own data and observe the effect of different parameter settings used in the chromatogram alignment. The DrawAlignR software uses the highly optimized C++ code to perform pairwise alignment, through which the effect of smoothing parameters, gap penalties, similarity type can be easily investigated. We envision it to be used not only for selecting optimum parameters for alignment, but also to verify imputations in peptide data-matrix through on-the-fly cross-run visualization of a precursor.

DrawAlignR provides a graphical interface to visualize the alignment of multiple LC-MS/MS runs using raw chromatographic alignment. The user can begin by setting their working directory to one that contains an mzml folder with.chrom.mzml files and an osw folder containing.osw files, using the “Set Working Directory” button shown in Figure 2*a*. To load in one or more chromatographic files containing XIC traces generated from a targeted proteomics or DIA experiment, use the “Choose a Chromatogram File” button, as well as a corresponding assay library used to generate the data, using the “Choose a Library File” button. Raw chromatogram data is cached in memory upon file upload, for fast subsequent access. Afterwards, an individual peptide from one of the uploaded chromatograms can then be searched for using the “Peptide Name” drop down list and its charge state set using the “Peptide Charge” button. The selected peptide is then interactively visualized across the selected runs and the “Number of Plots” parameter can be adjusted to select which runs should be displayed on the right panel.

**Figure 2:**
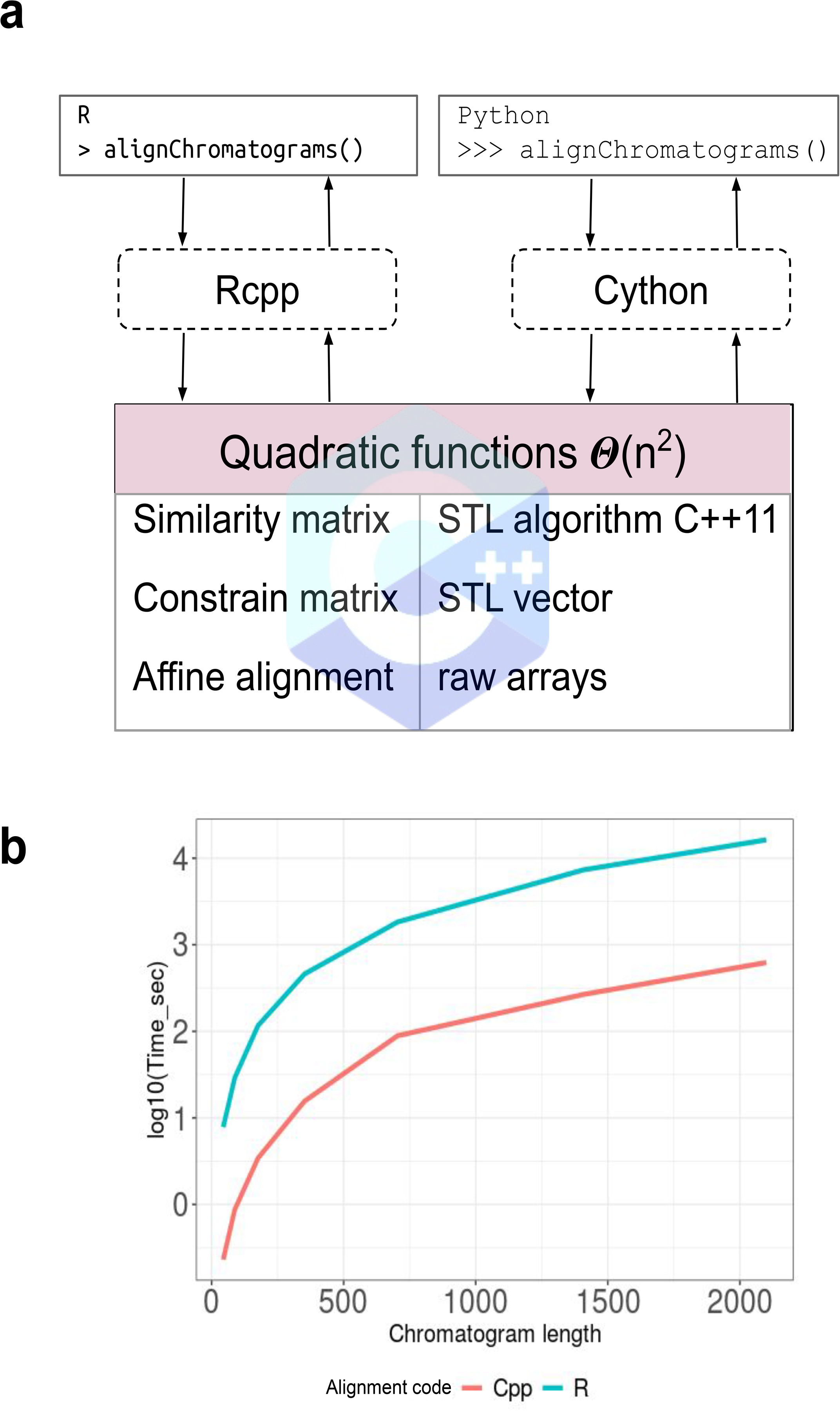
Screenshot demonstrating DrawAlignR interface and example case of visualization of DIAlignR’s alignment of two runs. a) shows the general settings options for setting the working directory, uploading chromatogram, library and osw result files, and choosing which runs to use as the reference or experiment if alignment is selected. The grey box in the second chromatogram plot depicts the zooming capability for further inspection of the data. b) shows all the alignment parameters that can be adjusted for testing and visualizing optimal alignment results.

If two or more chromatogram files are uploaded, the user can visualize alignment by first setting the Reference Run for Alignment and Experiment to Align parameters, and then checking the Plot Aligned button on the left panel, shown in Figure 2*a*. All available runs will then be aligned against the reference run and the user can compare pre and post-alignment chromatograms directly, using the “Reference Plot”, “Experiment Plot” and “Experiment Aligned Plot” buttons, thus observing the effect of the alignment interactively. The “Alignment Settings” panel also allows the user to change alignment parameters in order to fine-tune the alignment algorithm to their dataset, as shown in Figure 2*b*.

DrawAlignR has been tested using a manually annotated *Streptococcus pyogenes* dataset [2, 12] as well as a dilution series phosphoproteomics dataset [13] that were both acquired using a SWATH-MS acquisition scheme. A screenshot of an example precursor GEANVELTPELAFK is depicted in Figure 2, demonstrating the reference run XIC for hroest_K120808_Strep10%PlasmaBiolRepl1_R03, the experimental run for hroest_K120809_Strep10%PlasmaBiolRepl2_R04, and the shifted retention time of the aligned experimental run. Analysis is very fast due to our optimizations described above, allowing users to see alignment in real time and interactively. Loading the data takes less than a second for small dataset (and up to a minute for large datasets). After loading the data, users can explore their data interactively. A tutorial on how to use DrawAlignR is available at https://github.com/Roestlab/DrawAlignR/tree/master/vignettes.

## Outlook

Improving the quantitative consistency and ensuring that peptide peaks are correctly mapped across LC-MS/MS runs is a crucial step in obtaining a robust and accurate quantitative proteomics matrix. We have presented a significant improvement in the speed of DIAlignR alignment algorithm which decreases computation time from hours to minutes and on the one hand allows large-scale data processing and now allows an efficient interactive visualization of aligned chromatograms using DrawAlignR. We envision that this tool will be useful for validating alignment across multiple runs, and provide insight which can be used to fine-tune the parameters of the algorithm. The platform-independent C++ libraries have bindings to several popular languages such as R and Python and can easily be integrated in existing DIA analysis platforms such as OpenMS, TRIC and Skyline [5].

Furthermore, due to DrawAlignR’s modularity and ease of integration, it makes it possible to further extend the tool’s capability for visualizing other interesting chromatographic research questions, such as post-translational modifications (PTMs). Subsequently, SWATH-MS is very well suited for delving further into the proteome, to investigate post-translational modifications, due to its ability to separate isoforms of the same parent mass using site-determining product ions. [13] However, there currently is no visualization tool for verifying peptidoform-peak mapping assignments using site-determining ions, however, DrawAlignR can be easily extended for visual inspection.

## Contributions

HLR designed the project, SG and HLR implemented the C++ and Python code. AM and JS implemented the R Shiny app. HLR, SG, AM and JS wrote the manuscript.

## Acknowledgements

We are thankful for Oliver Alka and Leon Xu for discussion on writing C++ library.

## Funding

SG was supported by Mitacs Globalink Research Award, Cecil Yip Doctoral Research Award and Peterborough K.M. HUNTER Charitable Foundation Graduate Award. H.L.R. is supported by the Canada Research Chair program (#950-232260). This project was supported by Genome Canada (grant #15411) and the Ontario Genomics Institute (OGI-164), the CIHR (FRN: 161426).

## Data availability

DrawAlignR is available at https://github.com/Roestlab/DrawAlignR. DIAlignR code is available at Bioconductor or directly from Github (including Python bindings) at https://github.com/Roestlab/DIAlignR.

The authors have declared no conflict of interest.

## References

1. Aebersold, R., Mann, M. Mass-spectrometric exploration of proteome structure and function. Nature 537, 347–355 (2016) doi:10.1038/nature19949

2. Röst, H., Rosenberger, G., Navarro, P. et al. OpenSWATH enables automated, targeted analysis of data-independent acquisition MS data. Nat Biotechnol 32, 219–223 (2014) doi:10.1038/nbt.2841

3. Röst H.L., Aebersold R., Schubert O.T. (2017) Automated SWATH Data Analysis Using Targeted Extraction of Ion Chromatograms. In: Comai L., Katz J., Mallick P. (eds) Proteomics. Methods in Molecular Biology, vol 1550. Humana Press, New York, NY

4. Röst, H. L., Liu, Y., D’Agostino, G., Zanella, M., Navarro, P., Rosenberger, G.,…Aebersold, R. (2016). TRIC: an automated alignment strategy for reproducible protein quantification in targeted proteomics. Nature Methods, 13(9), 777–783.https://doi.org/10.1038/nmeth.3954

5. Egertson, J.D., MacLean, B., Johnson, R., Xuan, Y. & MacCoss, M.J. Multiplexed peptide analysis using data-independent acquisition and Skyline. Nat. Protoc. 10, 887–903 (2015).

6. Röst, H. L., Rosenberger, G., Aebersold, R., Malmstrom, L., Efficient visualization of high-throughput targeted proteomics experiments: TAPIR. Bioinformatics 2015, 31, 2415–2417.

7. Gupta S., Ahadi S., Zhou W., Röst H. DIAlignR provides precise retention time alignment across distant runs in DIA and targeted proteomics. Molecular & Cellular Proteomics 18, 806–817 (2019).

8. Wickham, H. (2019). Mastering Shiny. [online] Mastering-shiny.org. Available at: https://mastering-shiny.org/index.html [Accessed 26 Dec. 2019].

9. Eddelbuettel D, François R (2011). “Rcpp: Seamless R and C++ Integration.” Journal of Statistical Software, 40(8), 1–18. URL http://www.jstatsoft.org/v40/i08/.

10. Eddelbuettel D, François R. Writing a package that uses Rcpp. March 16, 2019, URL http://CRAN.R-Project.org/package=Rcpp.

11. Röst, H.L., Schmitt, U., Aebersold, R. & Malmström, L. pyOpenMS: a Python-based interface to the OpenMS mass-spectrometry algorithm library. Proteomics 14, 74–77 (2014).

12. Röst, H., Liu, Y., D’Agostino, G. et al. TRIC: an automated alignment strategy for reproducible protein quantification in targeted proteomics. Nat Methods 13, 777–783 (2016) doi:10.1038/nmeth.3954

13. Rosenberger, G., Liu, Y., Röst, H. L., Ludwig, C., Buil, A., Bensimon, A.,…Aebersold, R. (2017). Inference and quantification of peptidoforms in large sample cohorts by SWATH-MS. Nature Biotechnology, 35(8), 781–788. https://doi.org/10.1038/nbt.3908

14. Searle, B.C., Pino, L.K., Egertson, J.D. et al. Chromatogram libraries improve peptide detection and quantification by data independent acquisition mass spectrometry. Nat Commun 9, 5128 (2018) doi:10.1038/s41467-018-07454-w

15. HL Röst, L Malmström, R Aebersold. Reproducible quantitative proteotype data matrices for systems biology Molecular biology of the cell 26 (22), 3926–3931

16. Sturm M., Kohlbacher O. TOPPView: an open-source viewer for mass spectrometry data. J Proteome Res. 2009 Jul;8(7):3760–3. doi:10.1021/pr900171m.

17. Röst, H., Sachsenberg, T., Aiche, S. et al. OpenMS: a flexible open-source software platform for mass spectrometry data analysis. Nat Methods 13, 741–748 (2016) doi:10.1038/nmeth.3959

18. Röst, H. Python in proteomics, PeerJ Preprints 7, e27736v1

19. Levitsky, LI., Klein, JA., Ivanov, MV., Gorshkov, MV. Pyteomics 4.0: Five Years of Development of a Python Proteomics Framework. J Proteome Res. 2019 Feb 1;18(2):709–714. doi:10.1021/acs.jproteome.8b00717. Epub 2019 Jan 8.

