## Supplemental text for "DrawAlignR: An interactive tool for across run chromatogram alignment visualization"

### Contents

1. DIAlignR C++ Implementation
    - Note 1. Comparing C++ and R Implementation
    - Note 2. Organization of C++ Library and Interface Files
2. DrawAlignR User Manual
    - Getting Started
    - Performing Alignment
    - Graphical Elements

### DIAlignR C++ Implementation

#### Note 1. Comparing performance of C++ and R implementation of affine alignment

```
clen <- 704
set.seed(1)
A.XICs <- lapply(1:6, function(i){
  intensity <- rnorm(clen, mean = 10, sd = 10)
  time <- seq(from = 4964.7, to = 4964.7 + (clen-1)*3.41, by = 3.41)
  df <- data.frame("time" = time, "intensity" = intensity)
  df
})
B.XICs <- lapply(1:6, function(i){
  intensity <- rnorm(clen, mean = 10, sd = 10)
  time <- seq(from = 4978.4, to = 4978.4 + (clen-1)*3.41, by = 3.41)
  df <- data.frame("time" = time, "intensity" = intensity)
  df
})
intensityListA <- lapply(A.XICs, `[[`, 2) # Extracting intensity values
intensityListB <- lapply(B.XICs, `[[`, 2) # Extracting intensity values
tAVec <- A.XICs[[1]][["time"]]
tBVec <- B.XICs[[1]][["time"]]
B1p <- 4972.7
B2p <- 4972.7 + (clen-2)* 3.41
adaptiveRT <- 3.5*10
noBeef <- ceiling(adaptiveRT/3.14)
```

#### DIAlignR (v1.1.1)

```
system.time(for(i in 1:1000)
{DIAlignR::alignChromatogramsCpp(intensityListA, intensityListB, alignType
= "hybrid", tAVec, tBVec, normalization = "mean", simType =
"cosineAngle")})
```

#### DIAlignR (v0.1.0)

```
XICs <- list(list(A.XICs), list(B.XICs))
system.time(for(i in 1:1000) {
  s <- DIAlignR::getSimilarityMatrix(XICs, pep =1, runA =1, runB = 2, type
= "dotProductMasked")
  gapPenalty <- DIAlignR::getGapPenalty(s, gapQuantile = 0.5, type =
"dotProductMasked")
  Alignobj <- DIAlignR::getAffineAlignObj(s, go = gapPenalty*0.125, ge =
gapPenalty*40)
})

df <- data.frame("len" = c(44, 88, 176, 352, 704, 1408, 2100), "v0.1.0" =
c(7.81, 29.17, 116.62, 459.83, 1831.59, 7337.7, 16313.3), "v0.99.8"=
c(0.23,0.875,3.437, 15.596, 88.83, 266.97, 623.5))
df <- tidyr::gather(df, version, time, -len)
ggplot(df, aes(x = len, y= time/60, col = version)) + geom_line(size = 1.5)
+ theme_bw() + theme(legend.position="bottom", axis.text =
element_text(size=20), axis.title=element_text(size=15),
legend.text=element_text(size=15)) + xlab("Chromatogram length") +
scale_y_sqrt(breaks=c(0, 10, 100, 250)) + ylab("Time (min)")

df <- data.frame("len" = c(44, 88, 176, 352, 704, 1408, 2100), "R" =
c(7.81, 29.17, 116.62, 459.83, 1831.59, 7337.7, 16313.3), "Cpp"=
c(0.23,0.875,3.437, 15.596, 88.83, 266.97, 623.5))
df <- tidyr::gather(df, version, time, -len)
ggplot(df, aes(x = len, y= log10(time), col = version)) + geom_line(size =
1.5) + theme_bw() + theme(legend.position="bottom", axis.text =
element_text(size=20), axis.title=element_text(size=15),
legend.text=element_text(size=15)) + xlab("Chromatogram length") +
ylab("log10(Time_sec)")
```

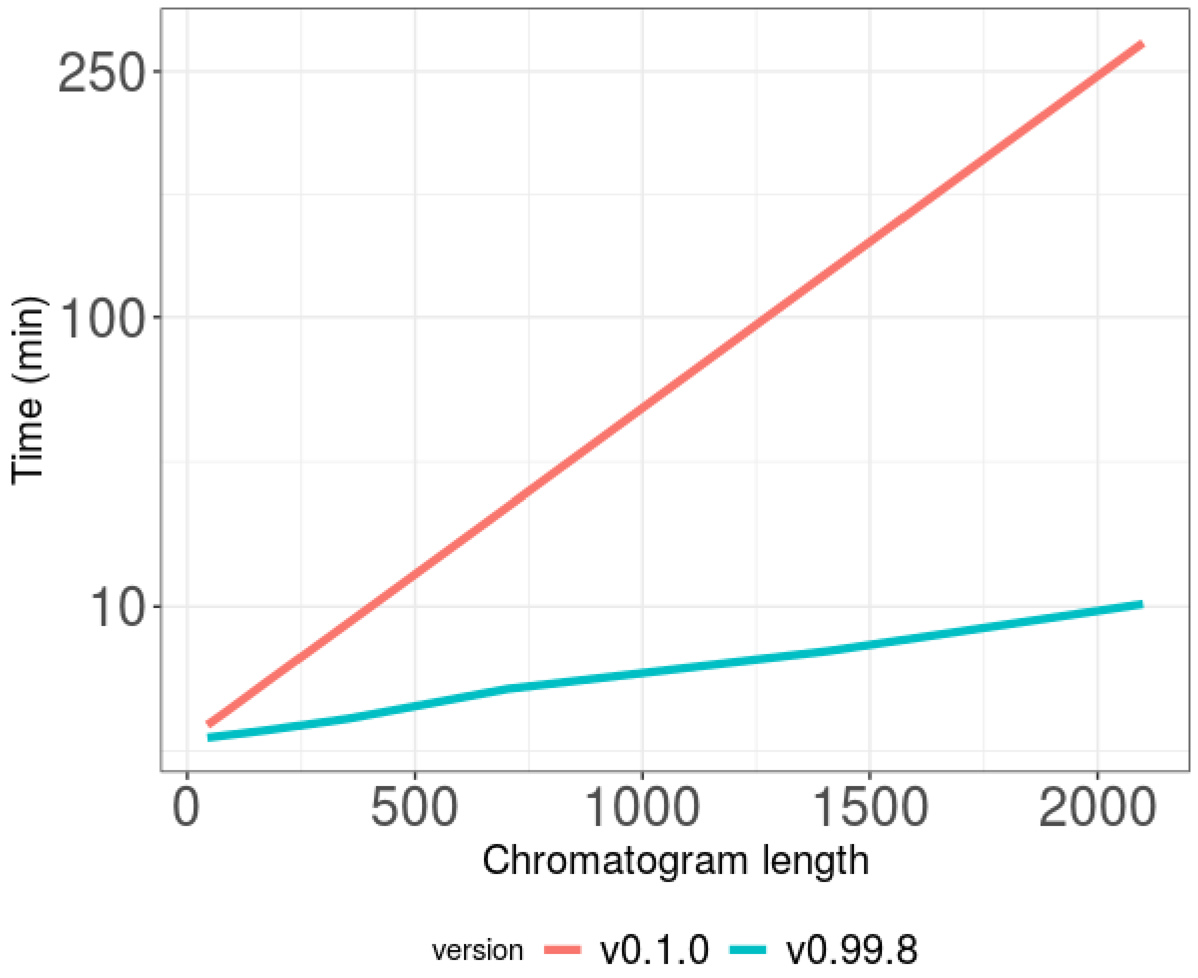


Quadratic function with respect to chromatogram length. DIAlignR version v0.1.0 has quadratic functions in R, whereas, latest version v0.99.8 has C++ implementation which provides faster execution than its predecessor. For each version, time taken by the algorithm to align 1000 peptides is plotted. Note the linear trend in execution time that is clearly observable, when the x-axis is plotted is in square root scale (as drawn here).

#### Note 2. Organization of C++ library and interface files to R in DIAlignR package

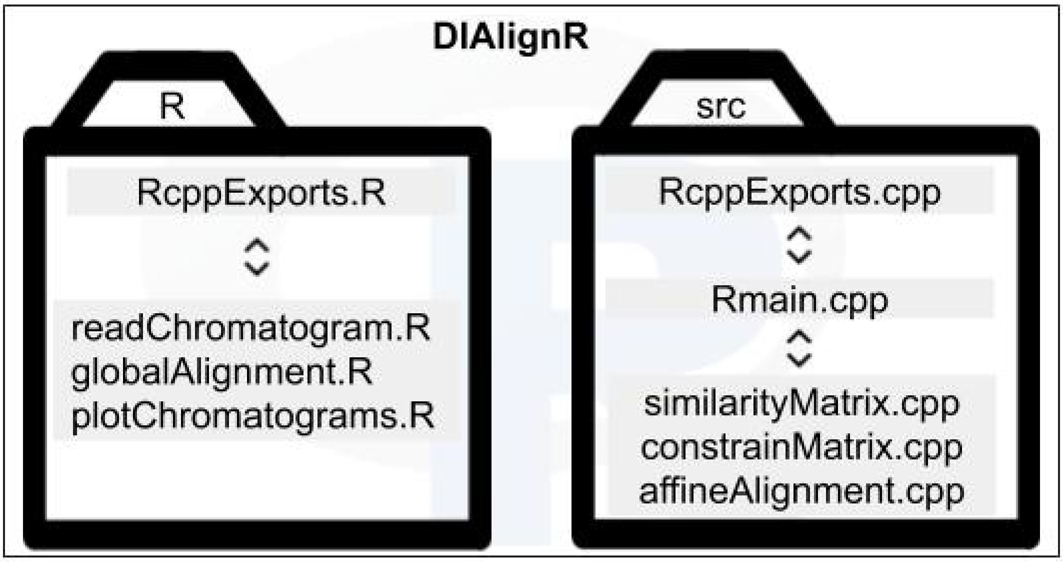

### DrawAlignR Tutorial

The visualization tool can be called via two methods, either via installing the repository using devtools github package installer or by calling shiny::runGitHub.
`Note:` You need to have R and RStudio installed.

#### Calling tool via Installing DrawAlignR package in R/ RStudio

```
### Install devtools if not already installed.
## install.packages("devtools")
## Install DrawAlignR from Roestlab Github Repository
devtools::install_github("Roestlab/DrawAlignR")
## To run the tool
DrawAlignR::runDrawAlignR()
```

OR

#### Calling tool via Github repo

```
### Install shiny
## install.packages("shiny")
shiny::runGitHub(repo = "Roestlab/DrawAlignR/", username = "Roestlab",
subdir = "inst/shiny-script")
```

### User Interface

There are three major tabs in the left side pannel:

- **General Settings**
  - These are general settings for uploading chromatogram files (.mzML or .sqMass), library assay files (.pqp), OpenSwathWorkflow results file (.osw). The user can also set the working directory that contains sub-directroys for osw and mzml files.
  - The user selects the peptide and charge state to visualize.
  - The user selects how many plots to show for each chromatogram run file suppled.
  - The user can select the alignment option to perform an alignment for the selected peptide.
  - The user can visualize the reference plot, experiment plot and the experiment aligned plot.
- **Alignment Settings**
  - The user can change various alignment parameters
- **Plot Settings**
  - The user can change various plot visualization settings

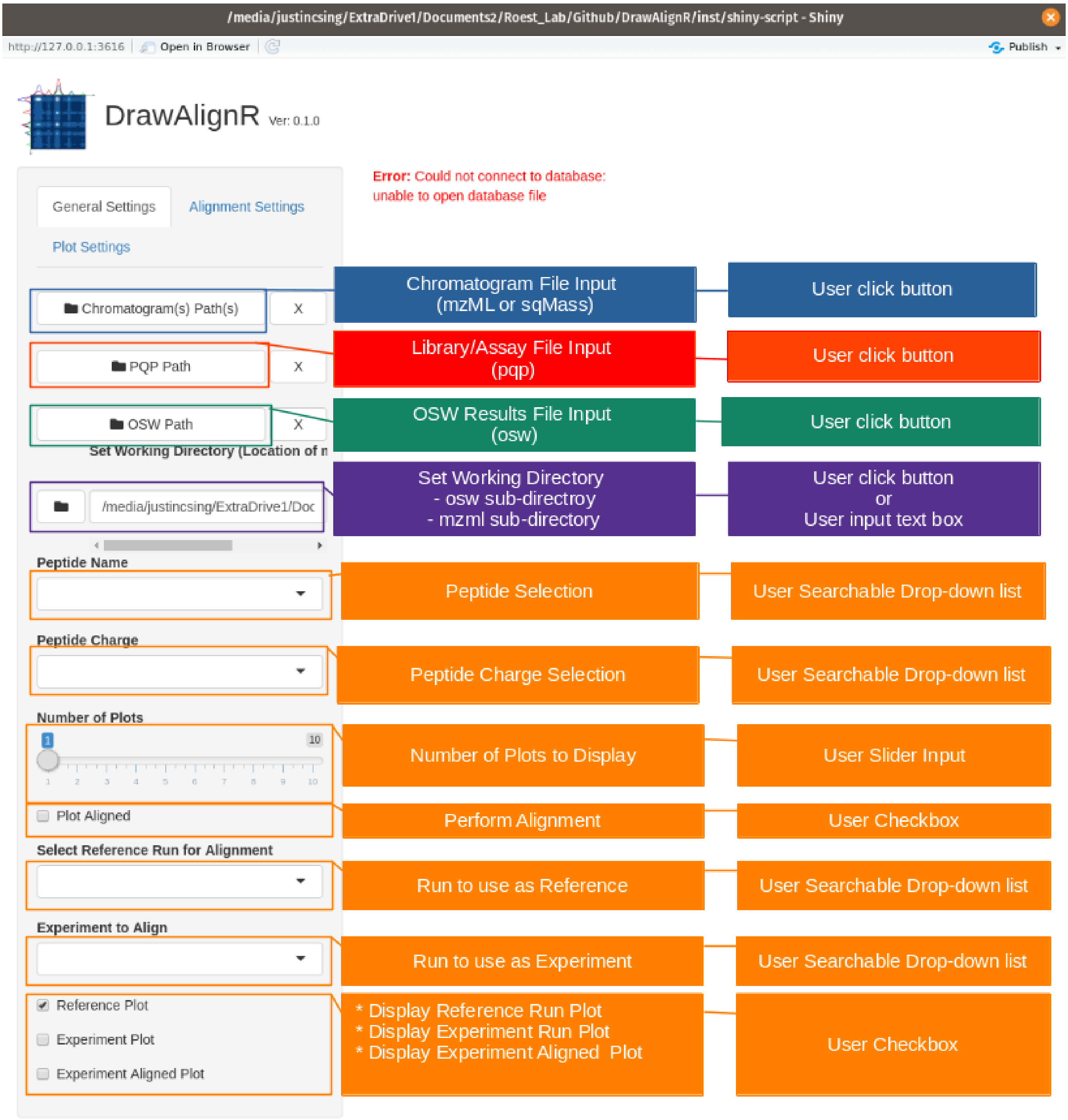
/media/justincsing/ExtraDrive1/Documents2/Roest_Lab/Github/DrawAlignR/inst/shiny-script - Shiny
http://127.0.0.1:3616
Open in Browser
Publish
DrawAlignR Ver: 0.1.0
Error: Could not connect to database: unable to open database file
General Settings
Alignment Settings
Plot Settings
Chromatogram(s) Path(s)
PQP Path
OSW Path
Set Working Directory (Location of n
/media/justincsing/ExtraDrive1/Doc
Peptide Name
Peptide Charge
Number of Plots
Plot Aligned
Select Reference Run for Alignment
Experiment to Align
Reference Plot
Experiment Plot
Experiment Aligned Plot
Chromatogram File Input (mzML or sqMass)
User click button
Library/Assay File Input (pqp)
User click button
OSW Results File Input (osw)
User click button
Set Working Directory
- osw sub-directroy
- mzml sub-directory
User click button or User input text box
Peptide Selection
User Searchable Drop-down list
Peptide Charge Selection
User Searchable Drop-down list
Number of Plots to Display
User Slider Input
Perform Alignment
User Checkbox
Run to use as Reference
User Searchable Drop-down list
Run to use as Experiment
User Searchable Drop-down list
* Display Reference Run Plot
* Display Experiment Run Plot
* Display Experiment Aligned Plot
User Checkbox

### Tutorial for Performing Alignment

#### Set Working Directory

Use the Set Working Directory button to set the working directory that contains an mzml folder with .chrom.mzml files and an osw file with a merged.osw file. Or you can directly enter the path to the working directory using the input textbox area.

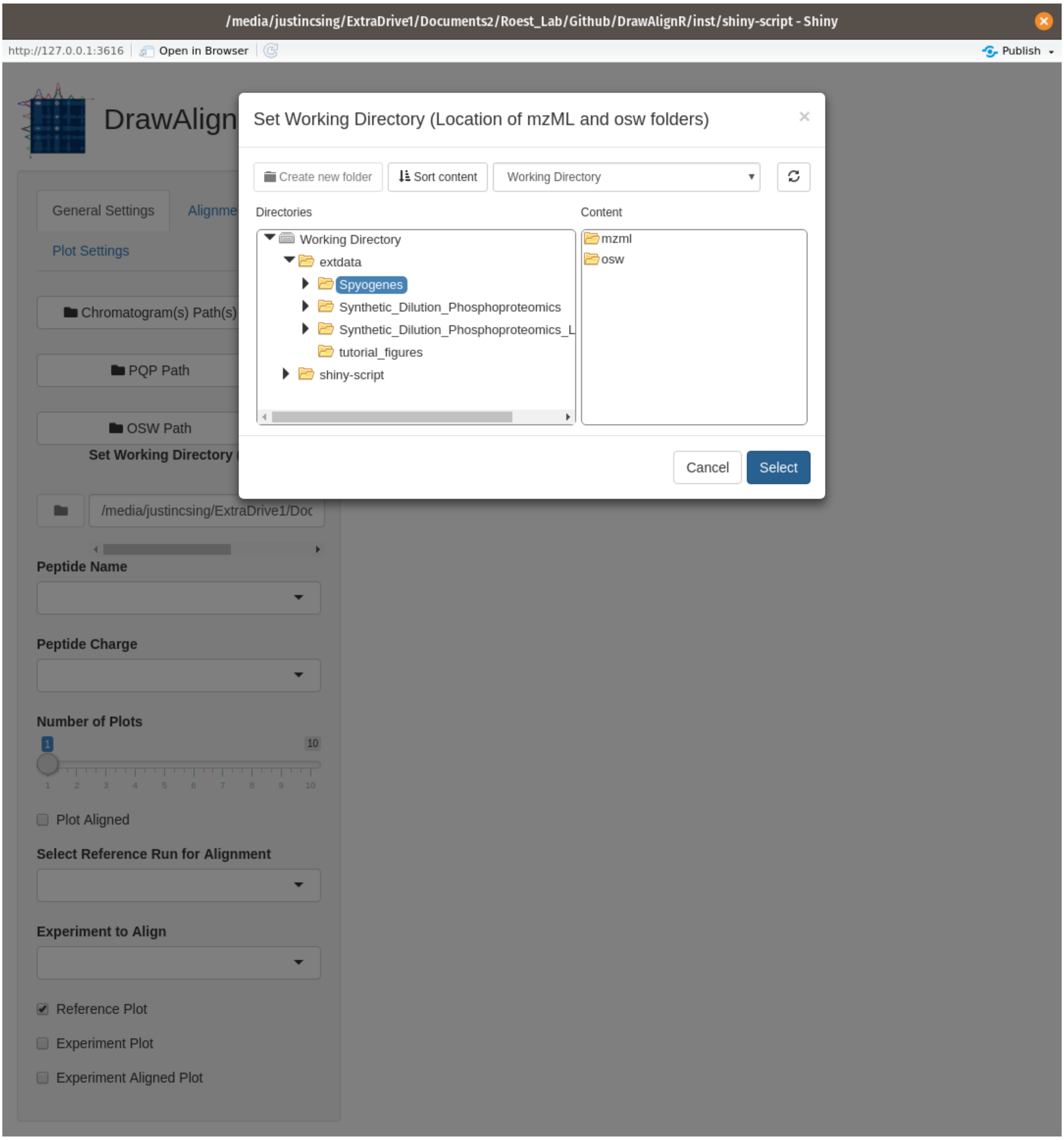

### Add Chromatogram files

Use the Choose a Chromatogram file button to select a chromatogram file(s) to upload. You can choose multiple files.

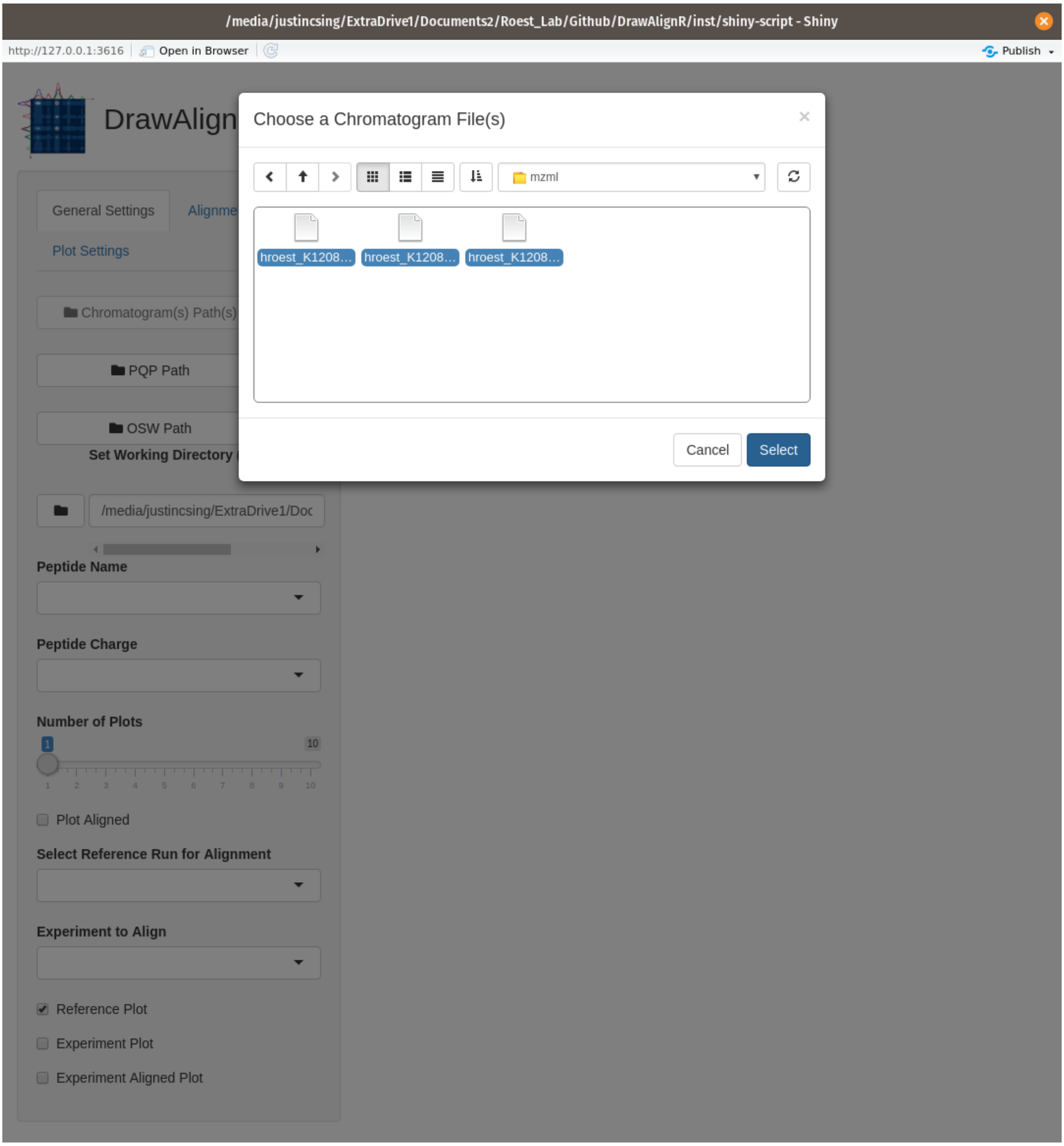

### Select a Peptide to visualize Chromatogram alignment

Choose which peptide you want to visualze using the Peptide dropdown list. The dropdown list is searchable, so you can easily search for a specific peptide to visualize. The list of peptides is extracted from either the input library file if abailable, or an osw file if available.

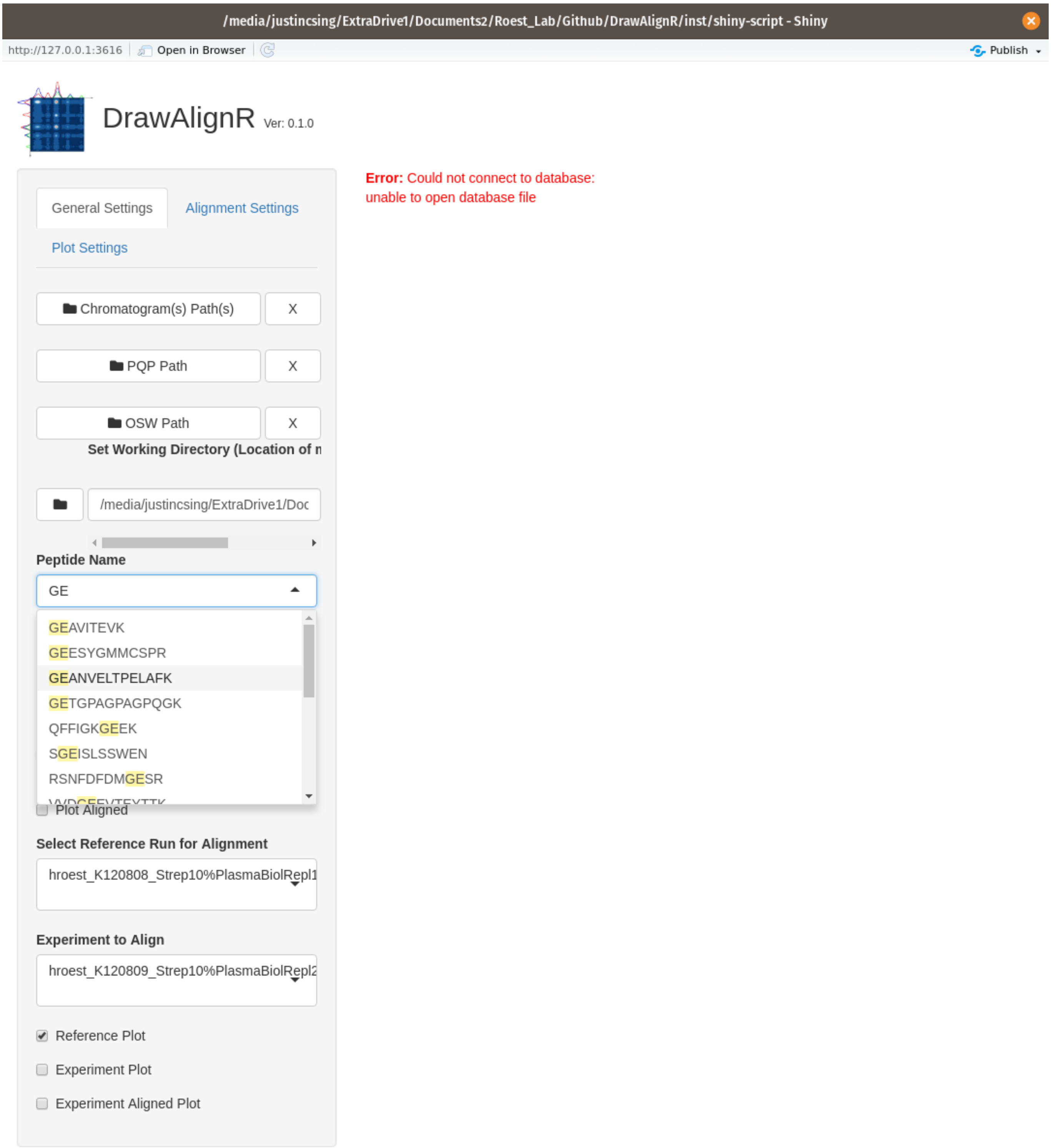

### Select a Charge State for Selected Peptide

Choose which charge state to visualize for the selcted peptide.

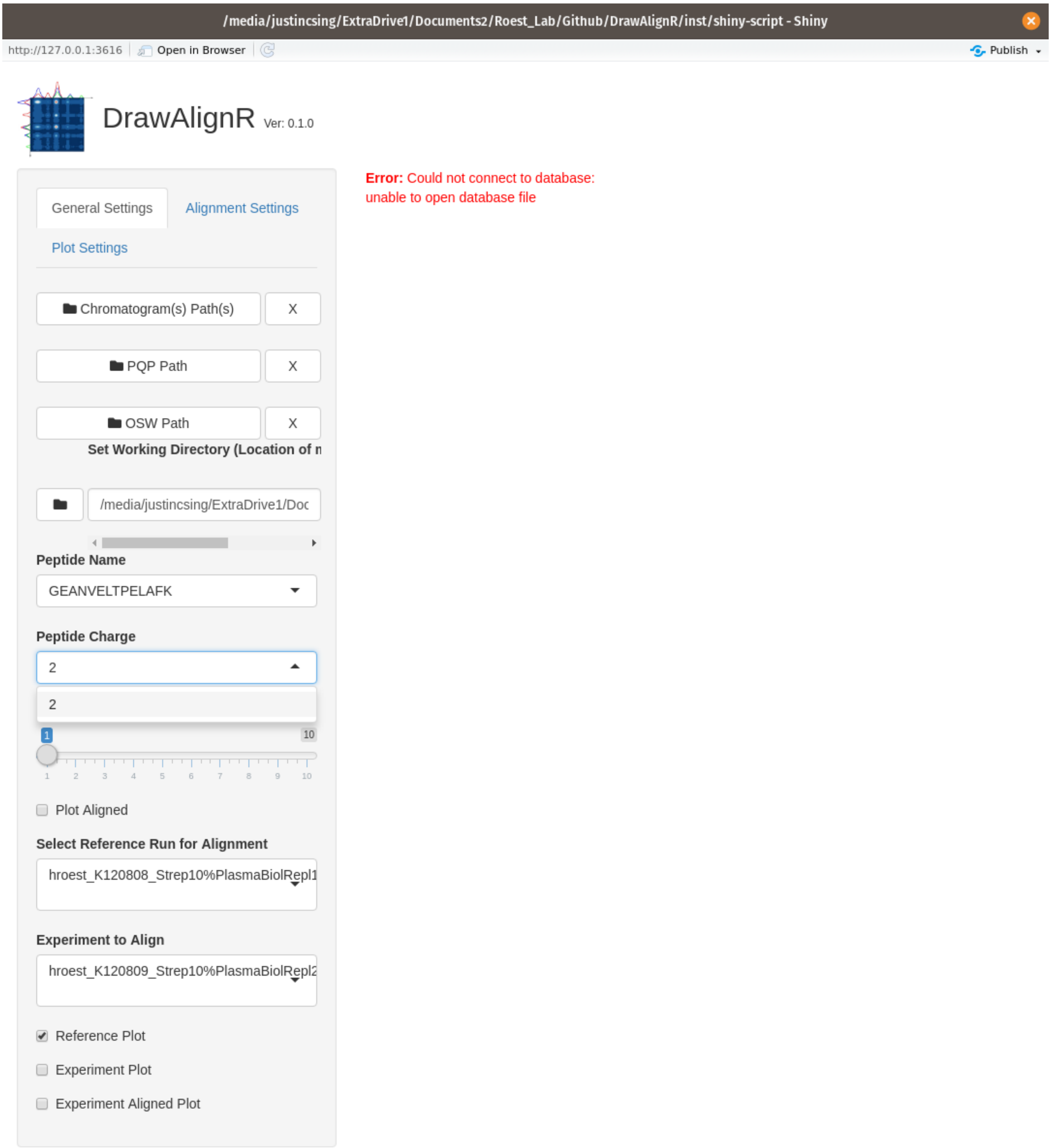

### Select Reference Run and Experiment Run

Choose which chromatogram file to use as the reference run, and which chromatogram file to use as the experiment run. These are set through the searchable dropdown lists, which extracts the filenames from the supplied chromatogram files without the .chrom.mzml extension

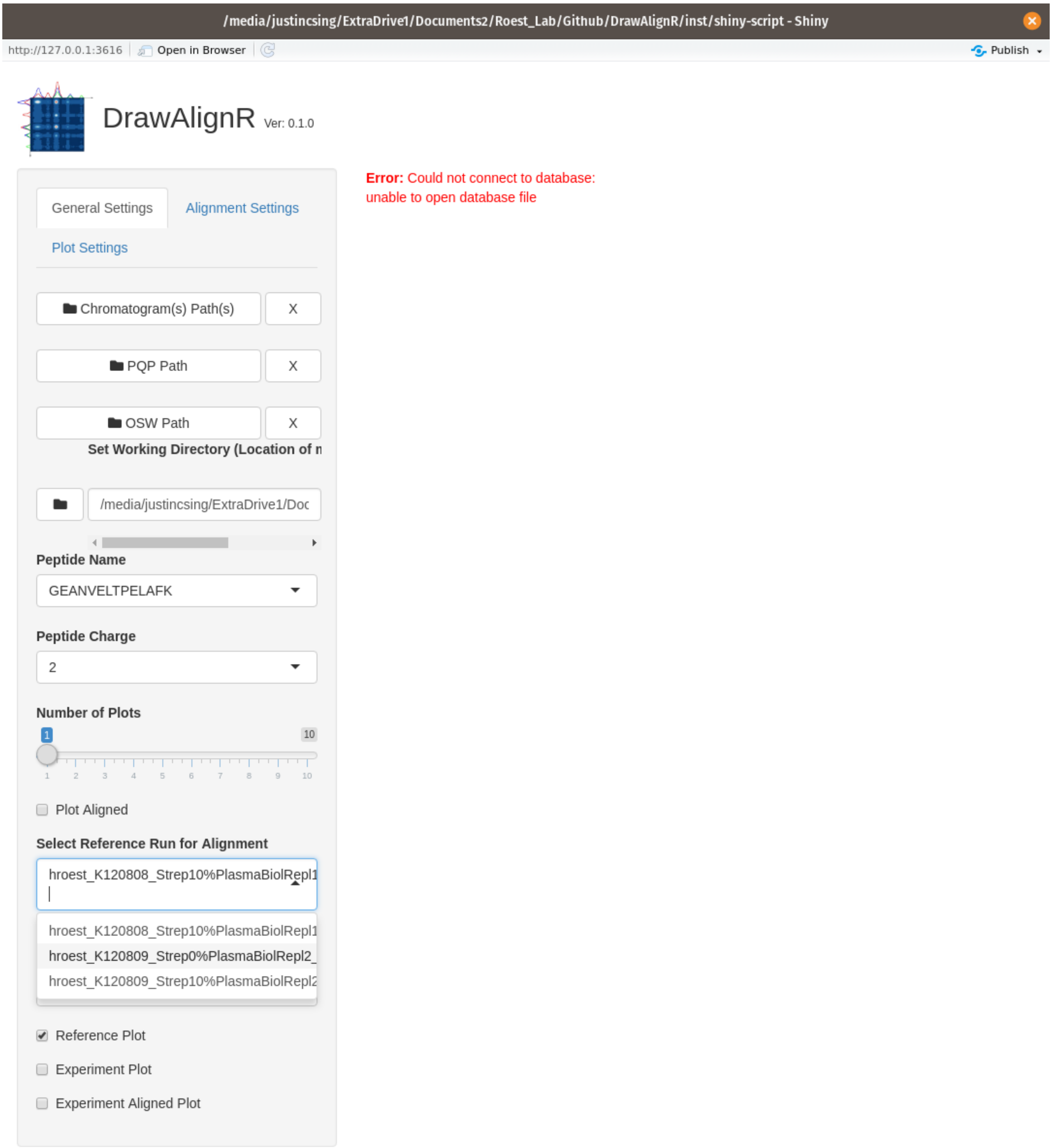

### Select Plot Align checkbox to perform alignment

Check the Plot Aligned checkbox to perform the alignment of the two runs for the seleected peptide You can also use the

- Reference Plot
- Experiment Plot
- Experiment Aligned Plot

to plot the different output extracted ion chromatogram results

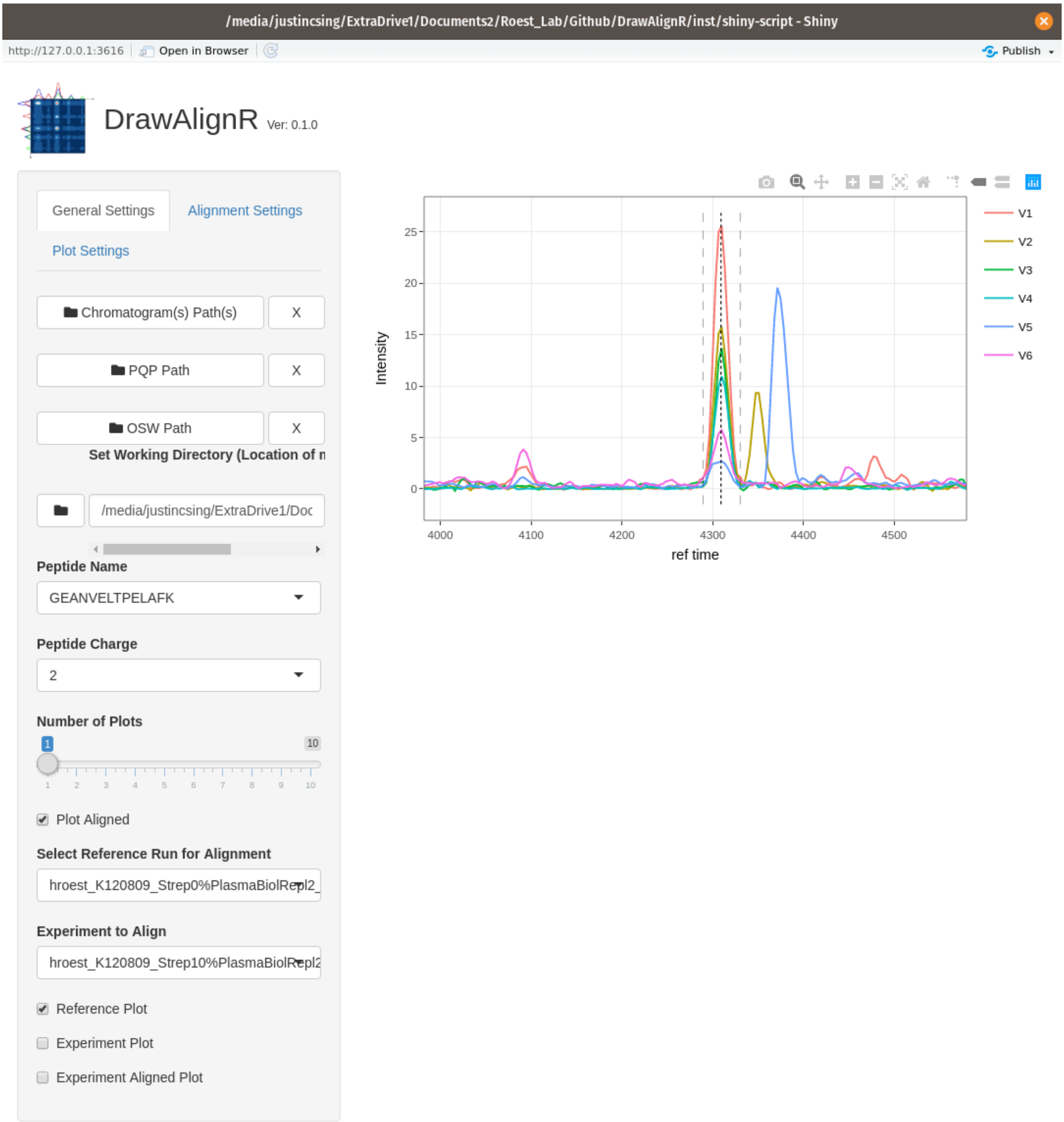

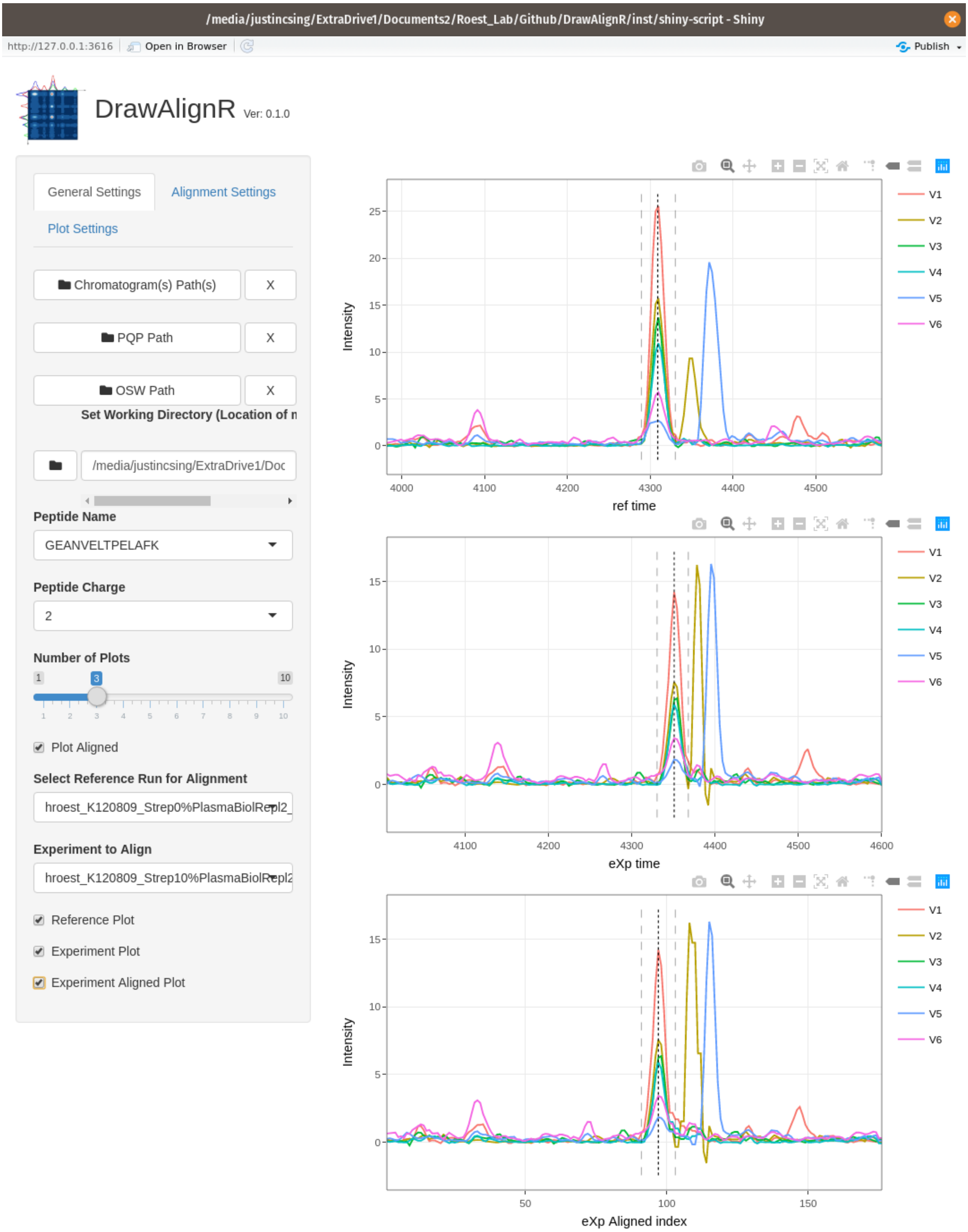
/media/justincsing/ExtraDrive1/Documents2/Roest_Lab/Github/DrawAlignR/inst/shiny-script - Shiny
http://127.0.0.1:3616
Open in Browser
Publish
DrawAlignR Ver: 0.1.0
General Settings
Alignment Settings
Plot Settings
Chromatogram(s) Path(s)
X
PQP Path
X
OSW Path
X
Set Working Directory (Location of n
/media/justincsing/ExtraDrive1/Doc
Peptide Name
GEANVELTPELAFK
Peptide Charge
2
Number of Plots
1
3
10
Plot Aligned
Select Reference Run for Alignment
hroest_K120809_Strep0%PlasmaBiolRepl2_
Experiment to Align
hroest_K120809_Strep10%PlasmaBiolRepl2
Reference Plot
Experiment Plot
Experiment Aligned Plot
Intensity
ref time
eXp time
eXp Aligned index
V1
V2
V3
V4
V5
V6

### Change Alignment Parameters

Select the Alignment settings tab to change various alignment settings

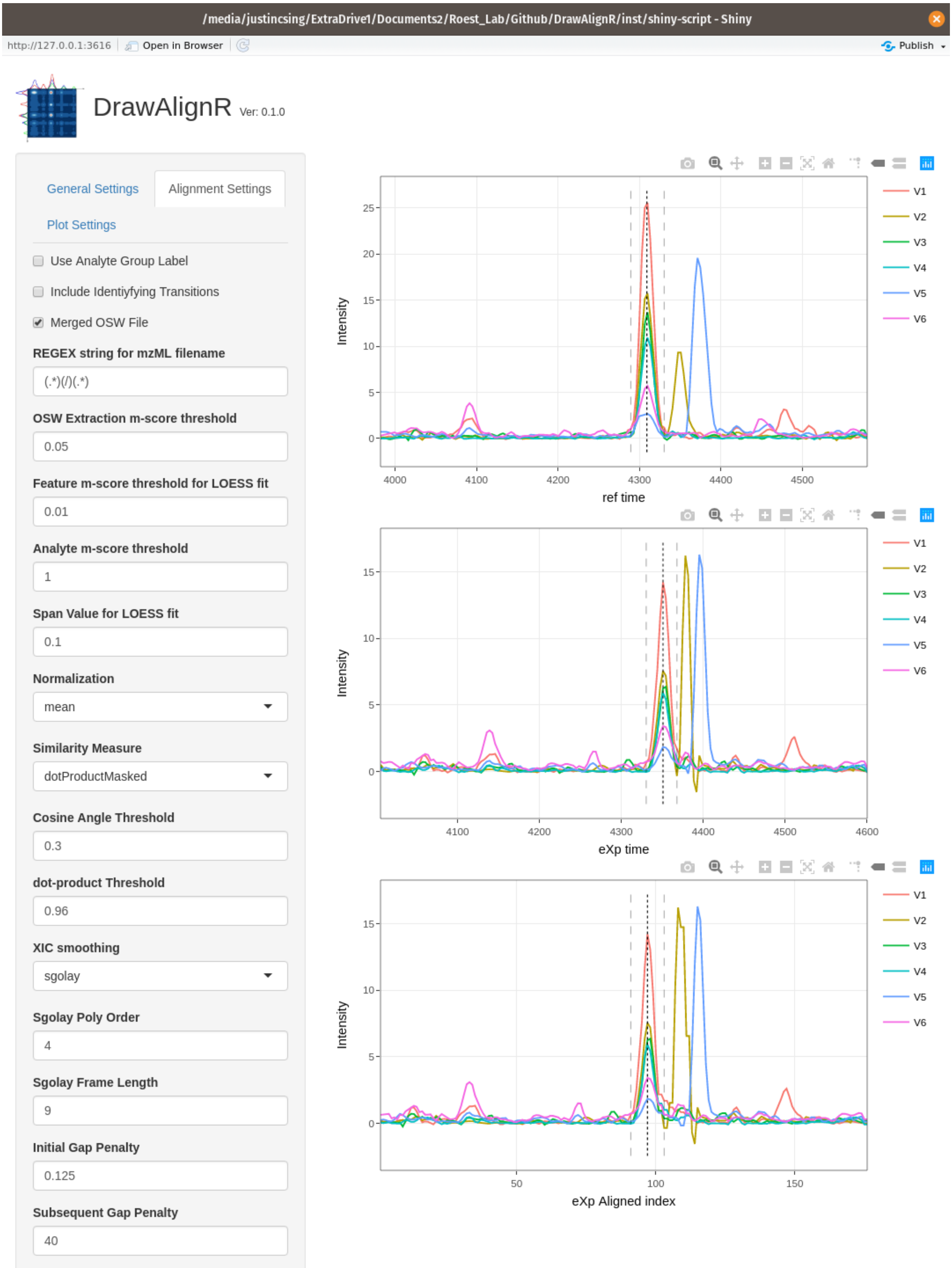

### Zoom into each chromatogram for further inspection

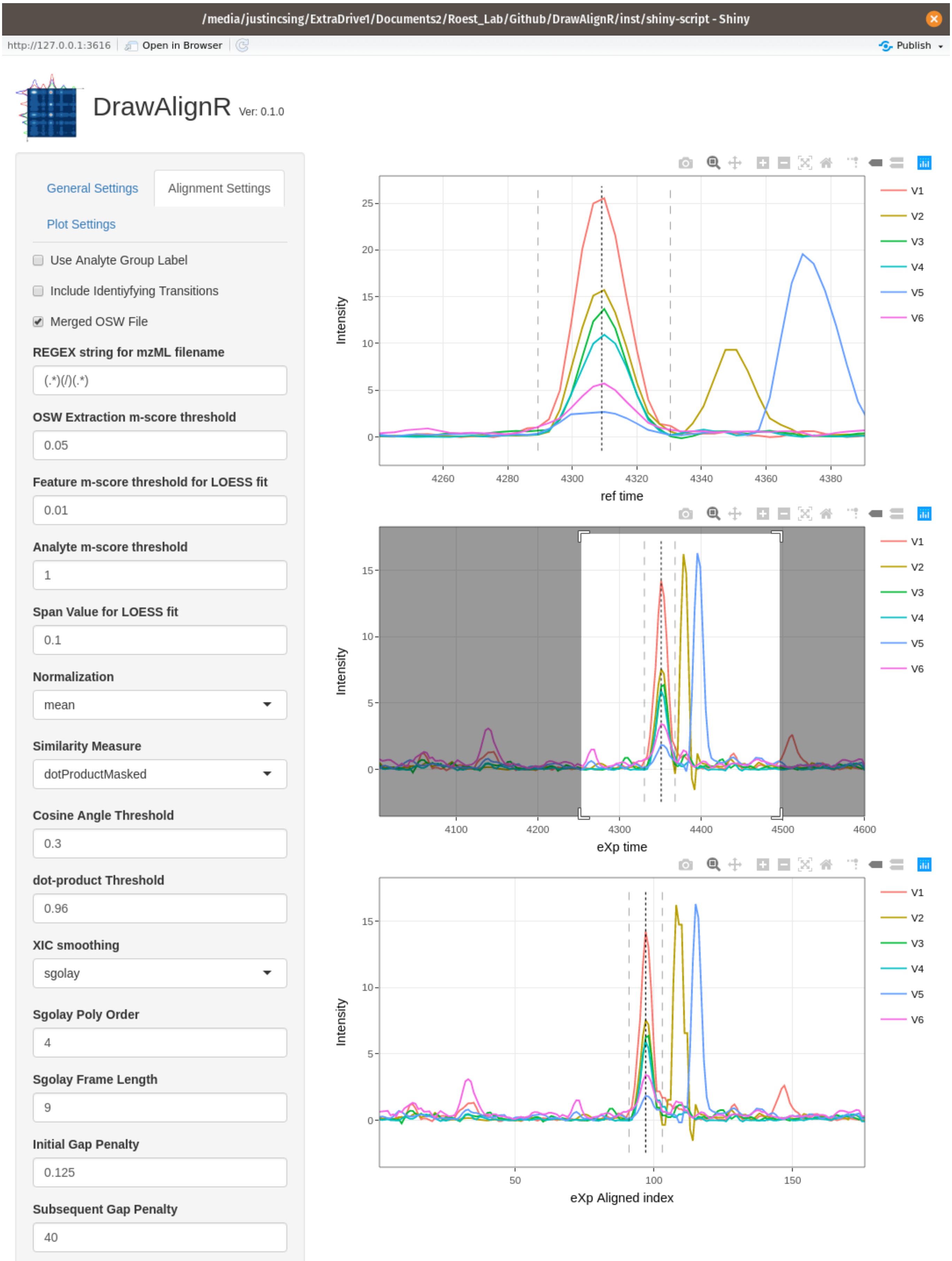

### Use the Hover-tooltip

You can hover over the chromatogram traces to see information such as Retention time and Intensity

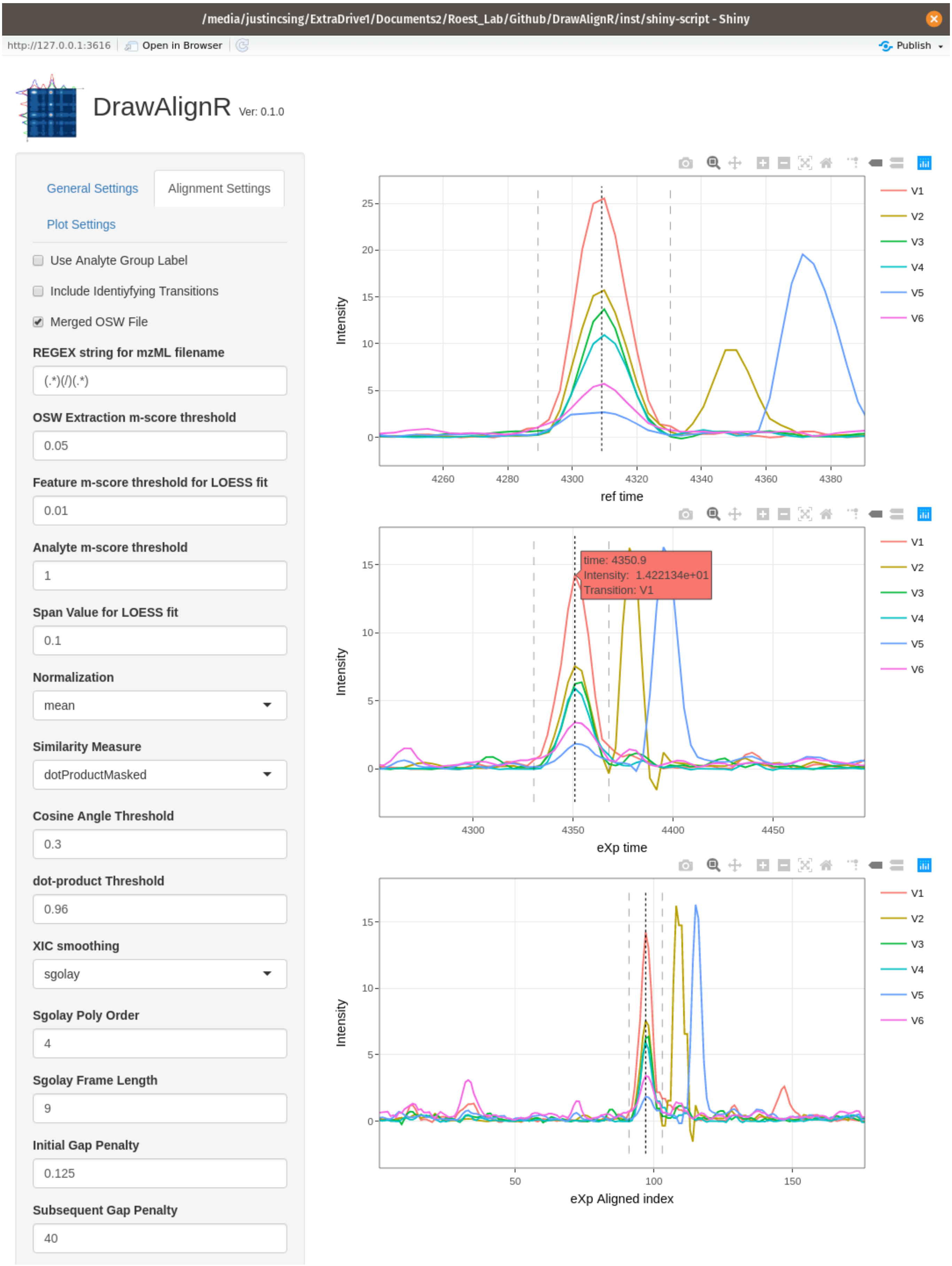

### Visualizing Chromatograms

The user can also just visualize the chromatograms alone without performing alignment to visually inspect each trace. If the user has an IPF dataset, they can visualize site-determining ions (unique identifying transitions) of the modified peptide.

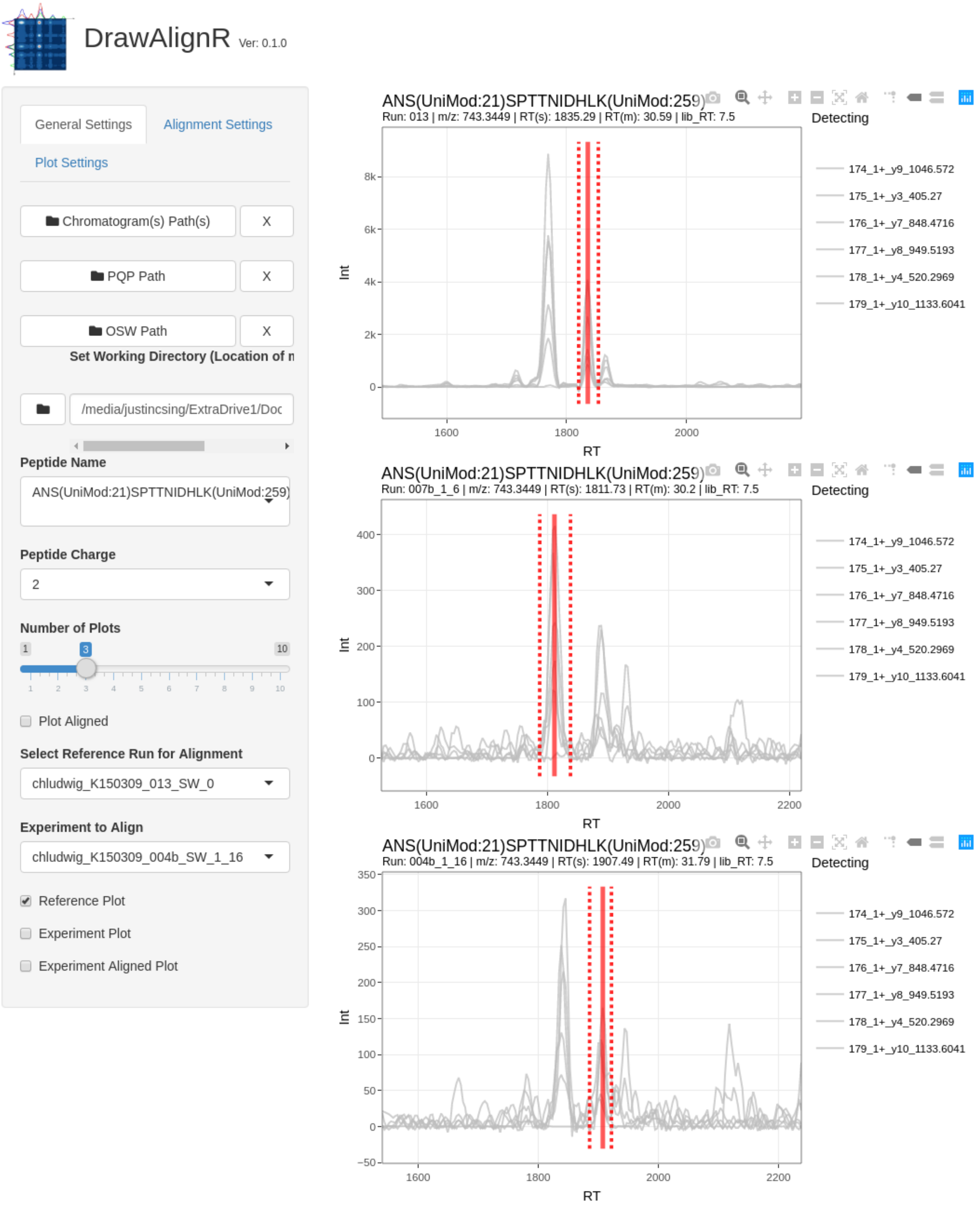

#### Precursor, Detecting, Identifying

The user can choose to display the precursor tace, or the 6 detecting traces, or the unique identifying traces. The precursor trace is displayed in `black`, the detecting traces are displayed in a `light gray` and the unique identifying transitions are `colored`

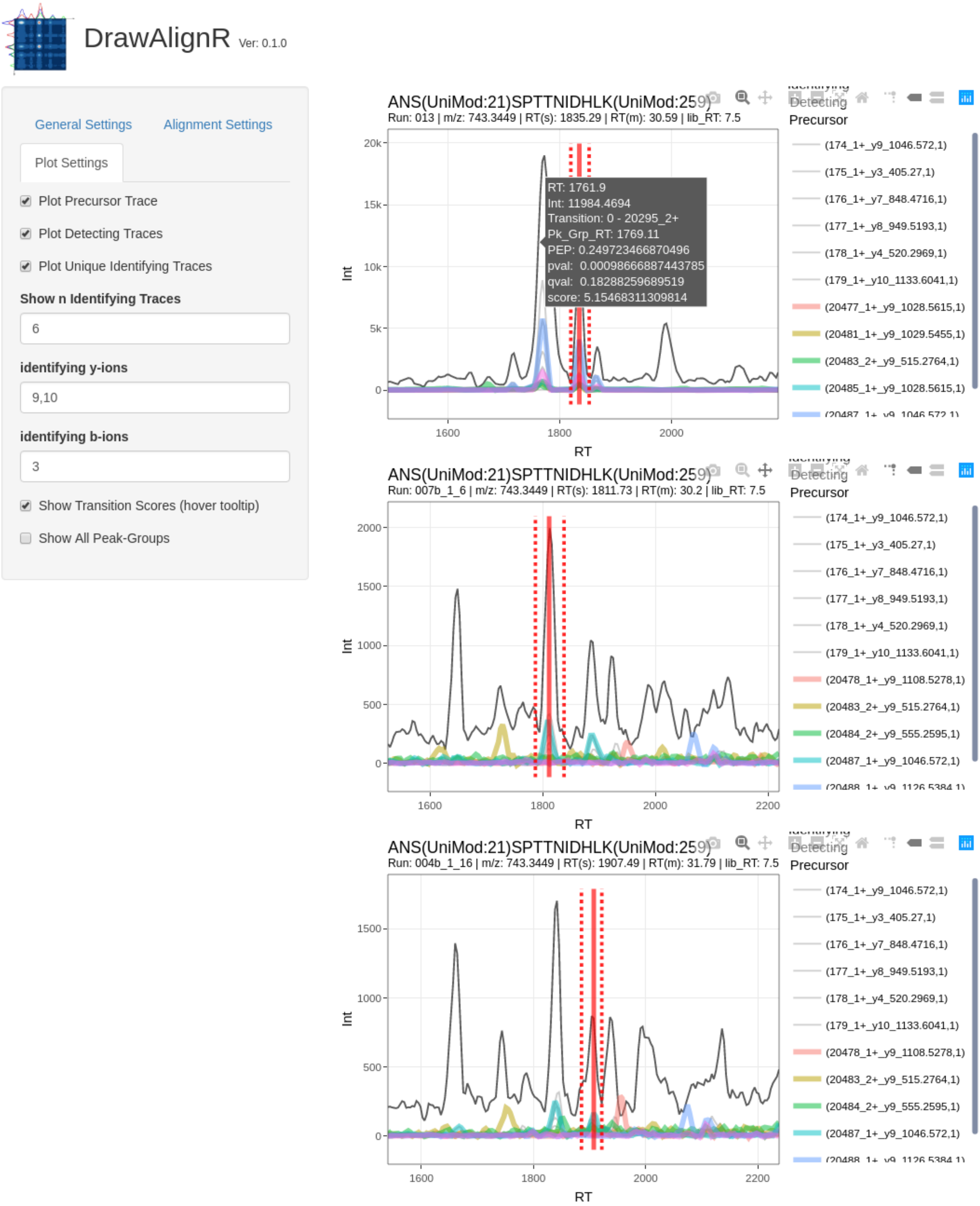

#### Displaying Transition Scores

The user can hover of the traces to display the transition scores such as the transitions posterior error probability, q-value and score.

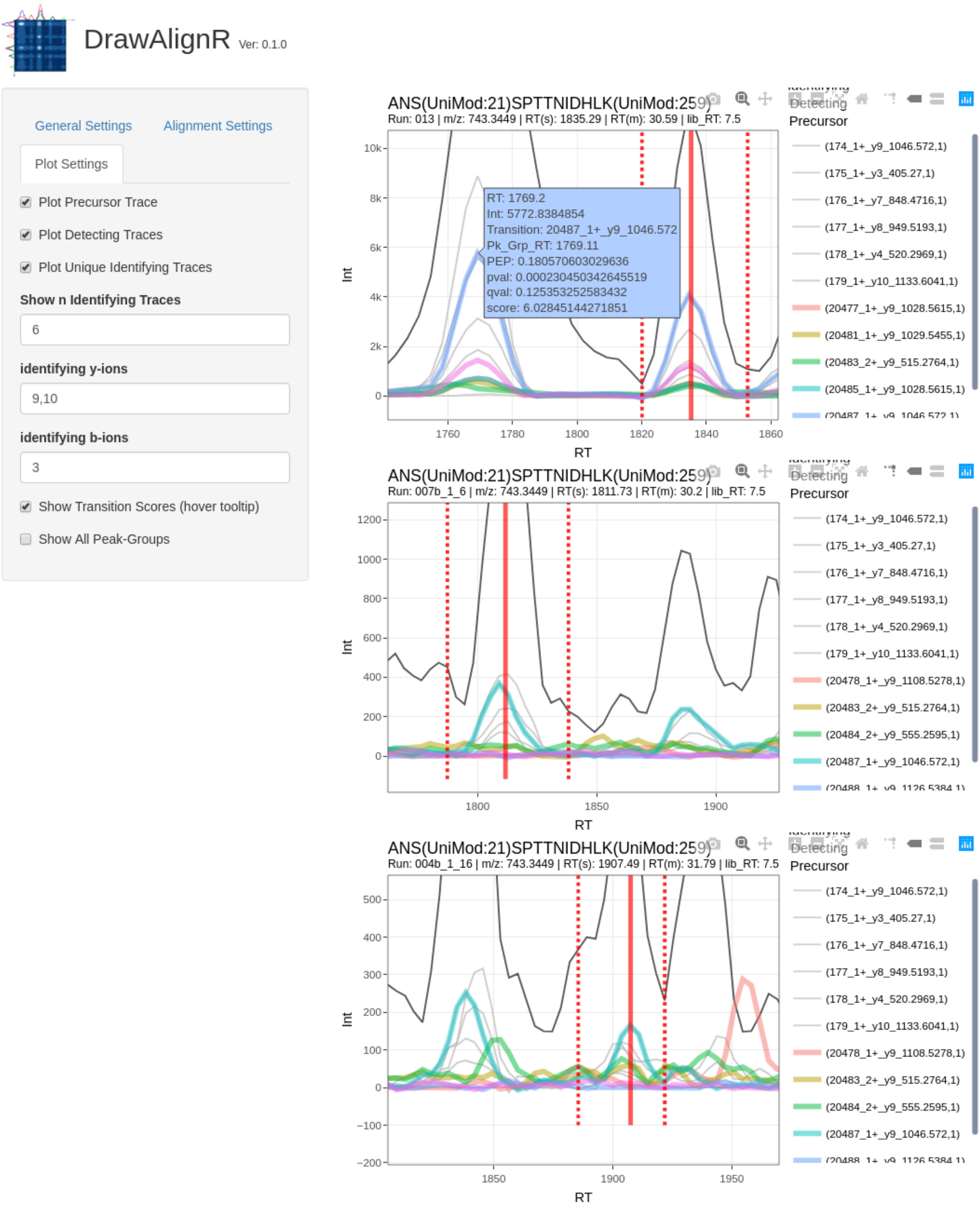

### Displaying all Peak-Group Ranks

The user can choose to display the other potential peak-group ranks found by OpenSWATH

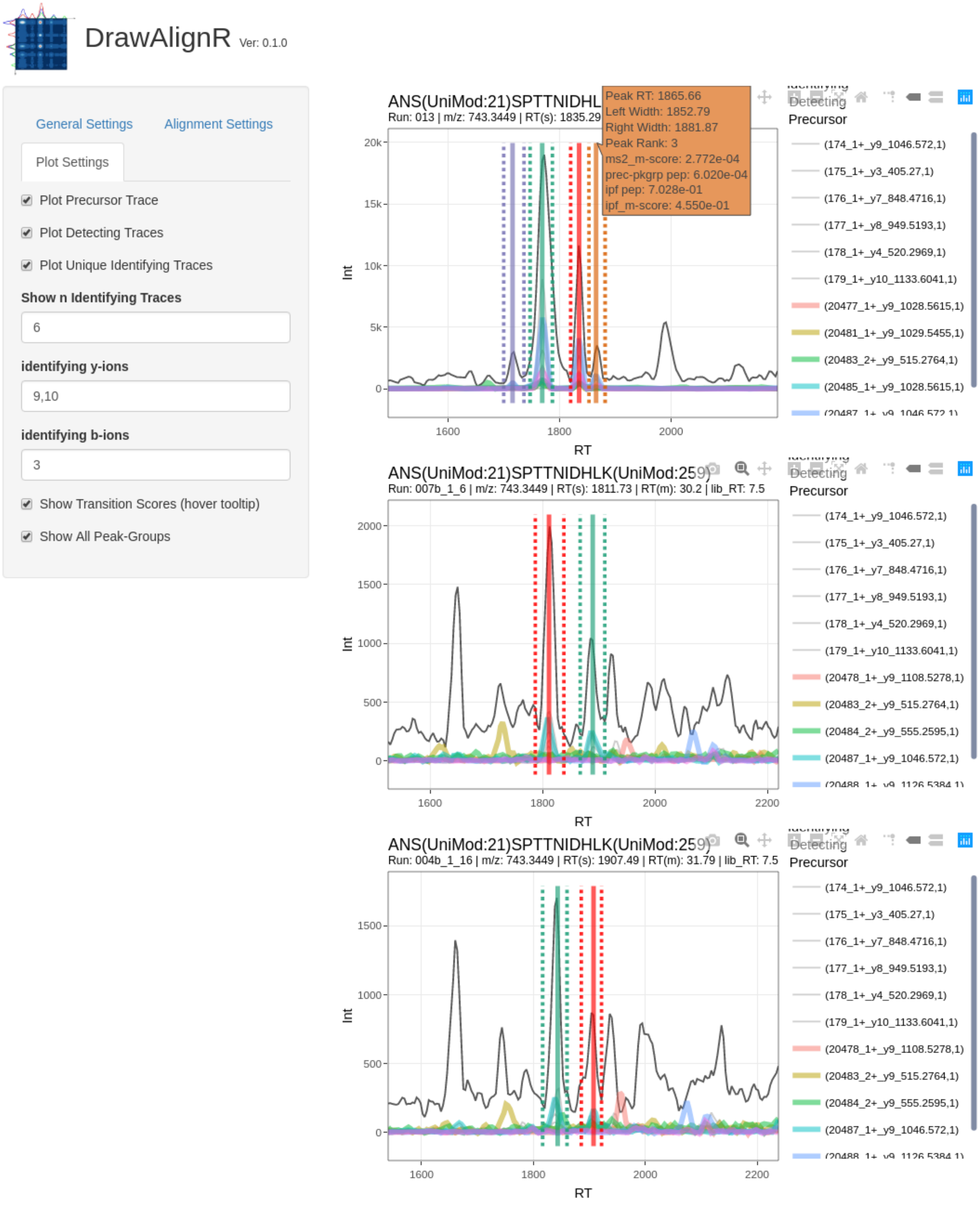

### Unselecting a Few Transitions to Display

The user can click on the legend to hide transitions they don't want to display

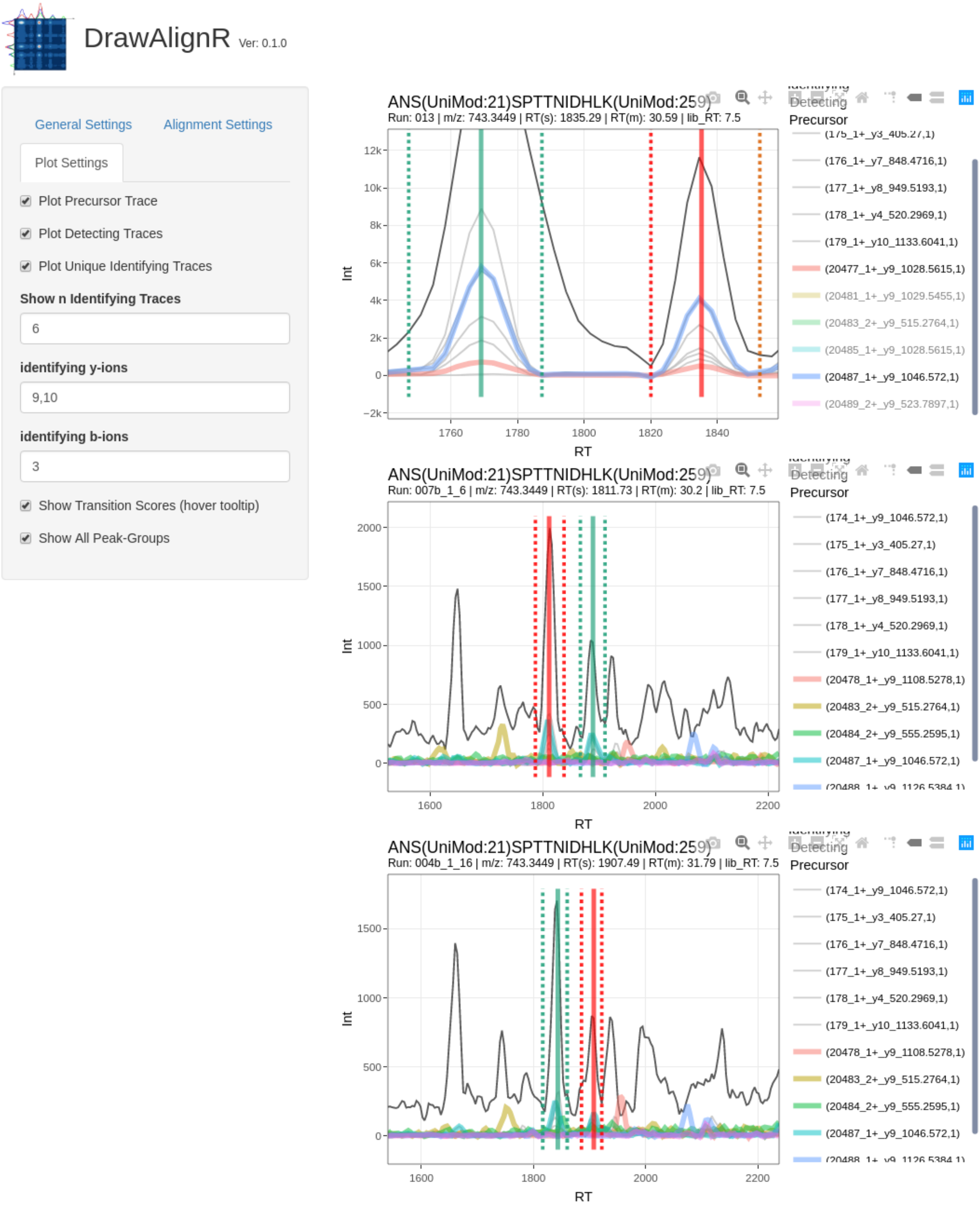

### Displaying a Single Transition

The user can choose to display a single transition by double clicking on the transition legend they wish to dispaly

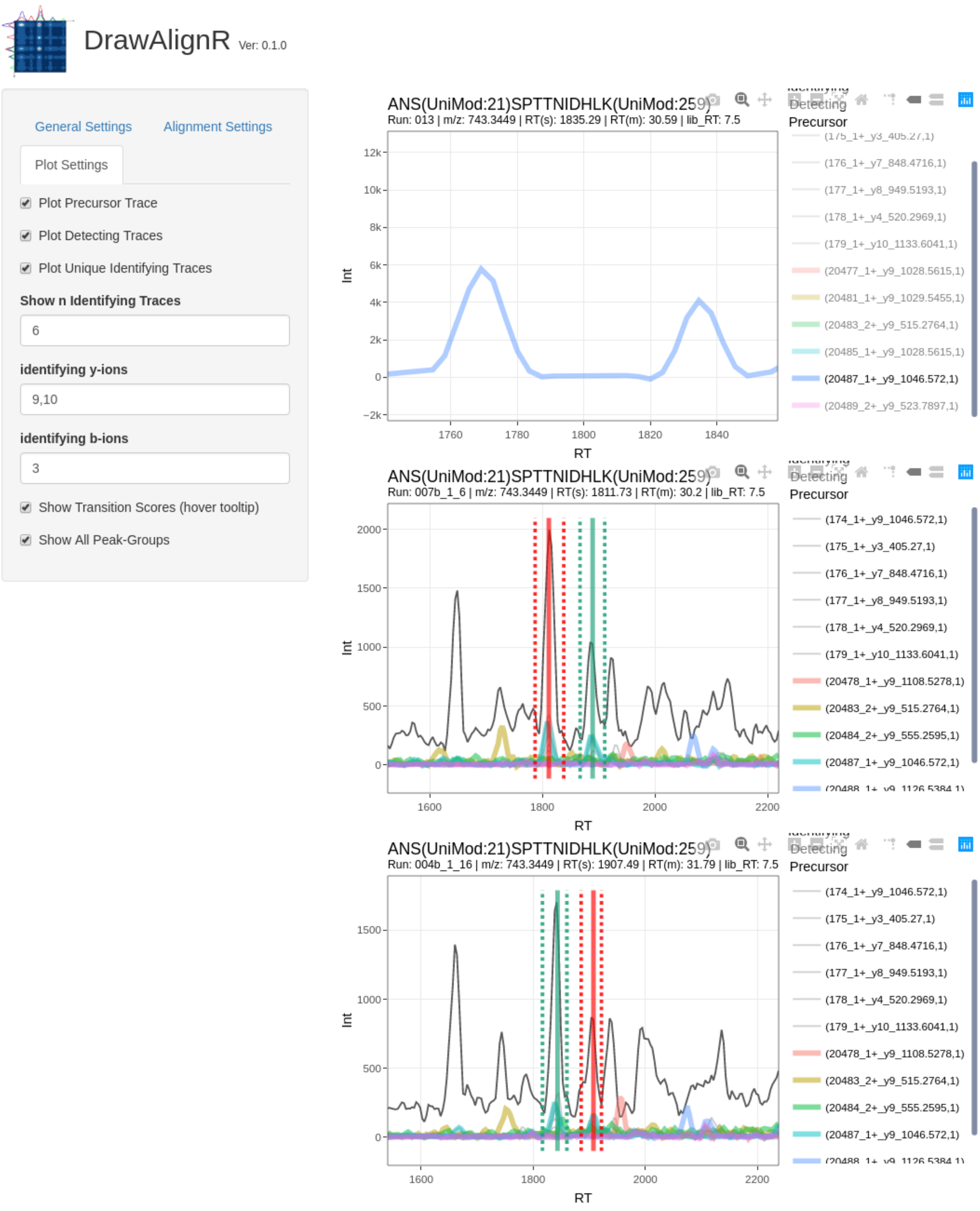
